# Transcriptional Control Of Calmodulin By CAMTA Regulates Neural Excitability

**DOI:** 10.1101/2020.09.14.296137

**Authors:** Thanh T. K. Vuong-Brender, Sean M. Flynn, Mario de Bono

## Abstract

Calmodulin (CaM) is the major calcium ion (Ca^2+^) sensor in many biological processes, regulating for example the CaM kinases, calcineurin, and many ion channels. CaM levels are limiting in cells compared to its myriad targets, but how CaM levels are controlled is poorly understood. We find that CaM abundance in the *C. elegans* and *Drosophila* nervous systems is controlled by the CaM-binding transcription activator, CAMTA. *C. elegans* CAMTA (CAMT-1), like its fly and mammalian orthologues, is expressed widely in the nervous system. *camt-1* mutants display pleiotropic behavioural defects and altered Ca^2+^ signaling in neurons. Using FACS-RNA Seq we profile multiple neural types in *camt-1* mutants and find all exhibit reduced CaM mRNA compared to controls. Supplementing CaM levels using a transgene rescues *camt-1* mutant phenotypes. Chromatin immunoprecipitation sequencing (ChIP-Seq) data show that CAMT-1 binds several sites in the calmodulin promoter. CRISPR-mediated deletion of these sites shows they redundantly regulate calmodulin expression. We also find that CaM can feed back to inhibit its own expression by a mechanism that depends on CaM binding sites on CAMT-1. This work uncovers a mechanism that can both activate and inhibit CaM expression in the nervous system, and controls Ca^2+^ homeostasis, neuronal excitability and behavior.

## Introduction

Calmodulin-binding transcription activators (CAMTAs) are a highly conserved family of Ca^2+^-regulated transcription factors^1^. In plants, CAMTAs are transcriptional effectors of Ca^2+^/CaM signaling in response to biotic/abiotic stress^2–6^. Mammals encode two CAMTA proteins that are enriched in heart and brain tissue^7^. Loss of CAMTA1 in the nervous system induces degeneration of cerebellar Purkinje cells, ataxia and defects in hippocampal-dependent memory formation^8,9^. A variety of neurological disorders, including intellectual disability, attention deficit hyperactivity disorder (ADHD), cerebellar ataxia, and reduced memory performance have been reported in individuals with lesions in the human *CAMTA1* gene^10–12^. Mechanistically however, little is known about the origin of these neuro-behavioural phenotypes. We show here that the *C. elegans* ortholog of CAMTA, CAMT-1, regulates neuronal Ca^2+^ signaling by controlling CaM expression. A variety of behaviours are dependent on CAMT-1, and Ca^2+^ imaging in multiple neurons reveals that neural activity is abnormal in *camt-1* mutants. By combining cell-type specific transcriptional profiling of these neurons with chromatin immunoprecipitation (ChIP) analysis of CAMT-1’s DNA-binding sites, we find that CAMT-1 upregulates the expression of CaM to control neural activity and behaviours. CaM is a ubiquitously expressed Ca^2+^ sensor that plays a key role in buffering intracellular Ca^2+^ and in directing a cellular response to Ca^2+^ changes^13,14^. It is pivotal to diverse processes, including metabolic homeostasis, protein folding, apoptosis, vesicular fusion, and control of neuronal excitability^15,16^, for example through regulation of CaM kinase II activity. Importantly, cellular levels of CaM are limiting, compared to the concentration of CaM binding proteins^17^, and changes in CaM levels are likely to impact Ca^2+^/CaM regulation of downstream targets^18^. How CaM expression is regulated in specific cell-types and contexts is, however, poorly characterized. Our results identify CAMTA as a CaM regulator in neurons, and suggest CAMT-1 can both promote CaM expression and repress it, in a feedback loop by which CaM can negatively control its own expression by binding CAMT-1.

## Results

### CAMT-1 functions in neurons to regulate multiple behaviours

We identified mutations in *camt-1*, the sole *C. elegans* CAMTA, in a forward genetic screen for mutations that disrupt *C. elegans* aggregation behavior^19^. This behavior, where animals form groups (Extended Data Fig. 1a), is closely linked to escape from ambient oxygen levels (21% O_2_)^20,21^. The screen used an N2 strain defective in the neuropeptide receptor NPR-1, *npr-1(ad609)*, that aggregates similarly to most natural isolates of *C. elegans;* the N2 lab strain does not aggregate due to an *npr-1* gain-of-function mutation^22^.

CAMT-1 has the characteristic domain architecture of CAMTAs^1^: a DNA binding domain (CG-1), an immunoglobulin-like fold (IPT/TIG) similar to those found in non-specific DNA-binding/dimerization domains of other transcription factors, ankyrin repeats (ANKs), a putative Ca^2+^-dependent CaM binding domain (CaMBD) and multiple IQ motifs that are thought to bind CaM in a Ca^2+^-independent manner (Fig. 1a, Extended Data Fig. 1b-c,^23,24^). CAMT-1 also has predicted nuclear localisation and nuclear export signals (NLS/NES, Fig. 1a). *camt-1* mutants showed defective responses to O_2_ stimuli. *npr-1* mutant animals move slowly in 7% O_2_ but show immediate and sustained arousal in 21% O_2_ (Fig. 1b). Two *camt-1* mutant strains isolated in the screen that introduced premature stop codons, *db894* (Q222*) and *db973* (R631*), were hyperactive in 7% O_2_, and showed a much smaller increase in speed when switched from 7% to 21% O_2_ (Fig. 1a-b, data not shown). A strain bearing a 451 amino acid deletion in *camt-1, ok515^25^*, recapitulated these defects (Fig. 1a-b). A fosmid transgene containing a wild-type (WT) copy of the *camt-1* genomic locus rescued *camt-1* mutant phenotypes, restoring fast movement at 21% O_2_, and quiescence (slow movement) at 7% O_2_ (Fig. 1c). These results indicate that CAMT-1 is required for *C. elegans* to respond appropriately to different O_2_ levels.

**Figure 1:**
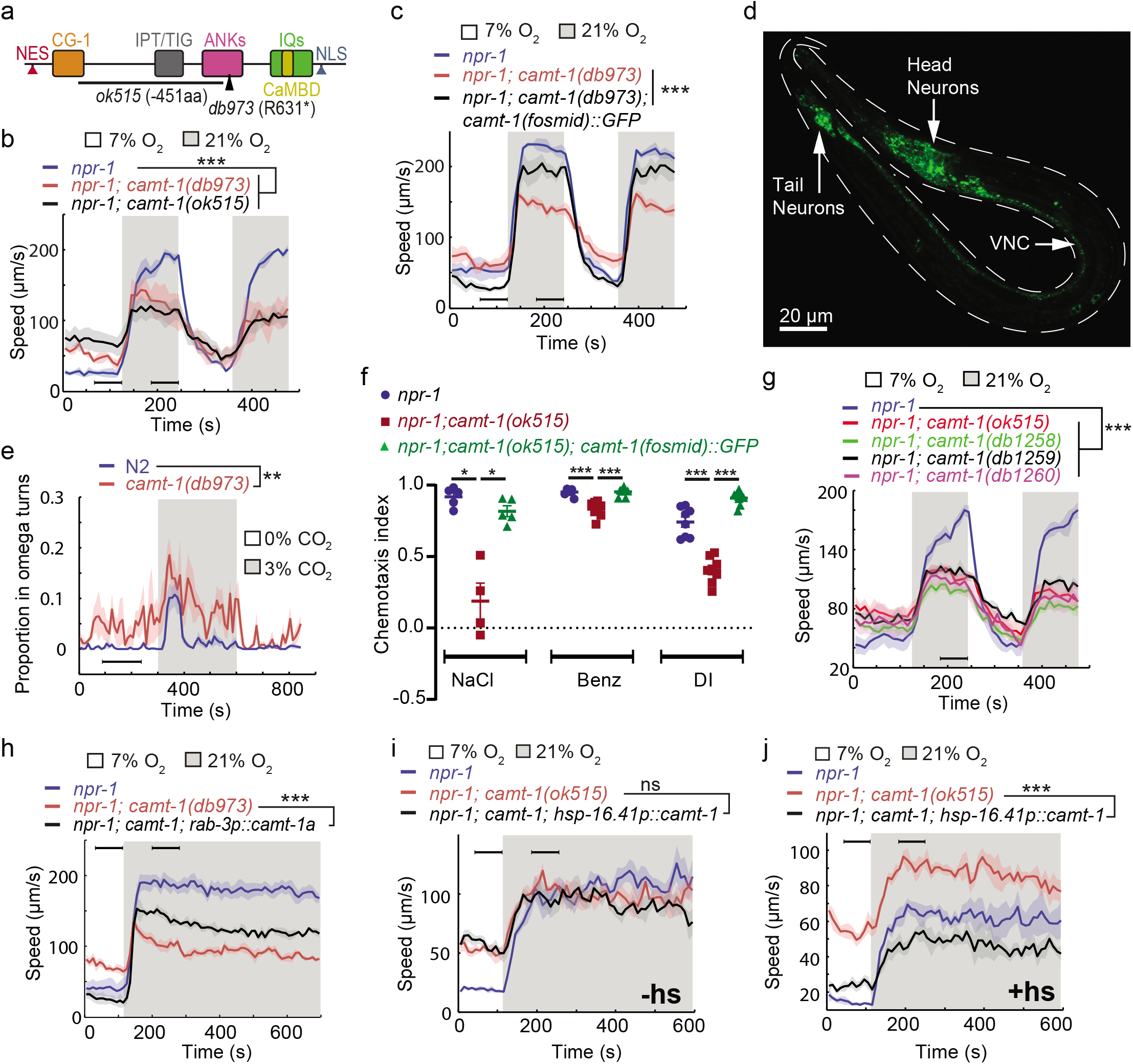
*camt-1* mutants exhibit pleiotropic behavioural defects. **a** The domain organization of CAMT-1, highlighting *camt-1* loss of function mutations used in this study. NES, nuclear export signal; CG-1, DNA-binding domain; IPT/TIG, Ig-like, plexins, transcription factors or transcription factor immunoglobulin; ANK, ankyrin domains; IQ, calmodulin binding motif; CaMBD, calmodulin-binding domain; NLS, nuclear localisation signal. **b** *camt-1* mutants exhibit altered locomotory responses to 21% O_2_ and hyperactive movement at 7% O_2_. **c** A WT copy of the *camt-1* genomic locus rescues the O_2_-response defects of *camt-1* mutants. **d** CAMT-1a::GFP driven from its endogenous regulatory sequences in a recombineered fosmid is expressed widely in the nervous system. VNC, ventral nerve cord. **e** *camt-1* mutants exhibit an increased turning frequency both in the presence and absence of a CO_2_ stimulus. Assays were performed in 7% O_2_. **f** *camt-1* mutants show defects in chemotaxis to NaCl, benzaldehyde (Benz) and diacetyl (DI), that can be rescued by expressing a WT copy of CAMT-1. Coloured bars indicate the mean and error bars the SEM. **g** The O_2_-response defects of mutants harbouring amino acid substitutions in the CG-1 DNA binding domain (*db1258, db1259* and *db1260* alleles; see Extended Data Fig.1b), are comparable to those of a *camt-1(ok515*) deletion mutant. **h** Pan-neuronal expression of the longest CAMT-1 isoform, CAMT-1a, in *camt-1* mutants, using the *rab-3* promoter, restores responsiveness to 21% O_2_ and low speed in 7% O_2_, although not fully. **i, j** Transgenic expression of CAMT-1 from the *hsp-16.41* heat shock promoter does not rescue the hyperactive locomotion of *camt-1(ok515)* mutants without heat-shock (**i**). Heat-shock induced expression of CAMT-1 in L4 animals rescues this phenotype in *camt-1(ok515)* mutants (**j**). **b**-**c, e** and **g**-**j**: Lines indicate average speed and shaded regions SEM. Black horizontal bars indicate time points used for statistical tests. *npr-1* denotes *npr-1(ad609)*. **b-c**, **e-j**: Mann-Whitney *U* test, ns: p≥ 0.05, *: p < 0.05, **: p < 0.01, ***: p < 0.001. Number of animals: n ≥ 22 (**b**), n>41 (**c**), n ≥ 23 (**e**), n≥4 assays for each genotype (**f**), n ≥ 56 (**g**), n ≥ 36 (**h**), n ≥ 42 (**i**), n ≥ 56 (**j**). hs: heat shock.

In mouse, humans and flies, CAMTA transcription factors are expressed in many brain regions^8–10,26^. We generated a fosmid-based reporter to map the expression pattern of the CAMT-1a, the longest isoform of *C. elegans* CAMTA. This reporter was functional, as it rescued the behavioural defects of *camt-1* mutants (Fig. 1c), and revealed that CAMT-1 is expressed broadly and specifically in the nervous system (Fig. 1d). We observed CAMT-1 expression in sensory neurons with exposed ciliated endings, motor neurons of the ventral cord, the URX O_2_-sensing neuron, and the RMG hub interneurons (Extended Data Fig. 2).

This broad expression prompted us to ask if *camt-1* mutants displayed pleiotropic behavioural phenotypes. We asked whether CAMT-1 is required for behavioural responses to chemical cues other than O_2_, and for other aversive behaviours, such as avoidance of CO_2_. In response to a rise in CO_2_, WT N2 worms transiently perform omega turns, Ω-shaped body bends that re-orient the animal away from the stimulus^27^. *camt-1* mutants exhibited abnormally high levels of omega-turns without a CO_2_ stimulus and a prolonged increase in omega turns in response to a rise in CO_2_ (Fig. 1e). *C. elegans* avoids CO_2_ but is attracted towards salt and a range of volatile compounds^28,29^. Chemotaxis towards NaCl, benzaldehyde and diacetyl was reduced in *camt-1* mutants, and these defects were rescued by a fosmid transgene containing WT CAMT-1 (Fig. 1f). Together these data show that CAMT-1 function is important for many *C. elegans* behaviours.

Many deleterious human alleles of CAMTA1 alter the CG-1 DNA binding domain^11^. We assessed the importance of the putative DNA-binding domain of CAMT-1 using mutants. Strains engineered using CRISPR to harbor mutations in conserved residues of the CG-1 domain showed defects in aggregation and in their response to O_2_, recapitulating phenotypes of the *camt-1* deletion mutants described above (Extended Data Fig.1b, Fig. 1g). By contrast, substituting conserved isoleucine residues in all four putative IQ domains to asparagines did not disrupt the O_2_-avoidance behaviors of *npr-1* mutant animals (Extended Data Fig. 1b and d). Moreover, expressing the a isoform of CAMT-1 with conserved residues in the CaMBD mutated rescued *camt-1* mutant phenotypes (Extended Data Fig. 1c and e). These results suggest that CAMT-1 binding to DNA is essential for its function but binding to CaM is not, at least for oxygen escape behaviour.

We targeted CAMT-1 cDNA expression to different subsets of neurons to find out where it is required to promote aerotaxis. Restricting expression of CAMT-1 to RMG but not O_2_ sensing neurons rescued the fast movement at 21% O_2_ in *camt-1* mutants (Extended Data Fig. 1f-g). Defective responses of *camt-1* mutants to 7% O_2_ was neither rescued by the expression of CAMT-1 in RMG nor simultaneous expression in RMG and O_2_-sensing neurons (Extended Data Fig. 1f-g). This data was consistent with a model in which CAMT-1 acts in multiple neurons. As expected, pan-neuronal expression rescued mutant phenotypes; in particular, expression of the a isoform alone (CAMT-1a) was sufficient for rescue (Fig. 1h). Interestingly, pan-neuronal overexpression of CAMT-1 reduced locomotory activity at 7% O_2_ to below levels found in the *npr-1* mutant background, suggesting that animal speed in 7% O_2_ is anti-correlated with CAMT-1 levels (Fig. 1h).

CaM/Ca^2+^-dependent changes in gene expression are known to be important for both development and function of the nervous system^30,31^. To test whether CAMT-1 activity is required during development, we expressed CAMT-1 cDNA from a heatshock-inducible promoter. Without heat-shock, this transgene did not rescue the hyperactivity phenotype of *camt-1* mutants (Fig. 1i). However, inducing CAMT-1 expression in late L4s/ young adults was sufficient to rescue the adult mutant defects (Fig. 1j), suggesting that CAMT-1 does not act developmentally to regulate behavioural responses to ambient O_2_.

### CAMT-1 dampens Ca^2+^ responses in sensory neurons

To test whether disrupting *camt-1* altered physiological responses to sensory cues we recorded stimulus-evoked Ca^2+^ changes in O_2_- and CO_2_-sensing neurons with Yellow Cameleon (YC) sensors. URX activity tracks environmental O_2_ levels, and tonic signaling from URX to RMG drives high locomotory activity at 21% O_2_^20^. BAG and AFD neurons are CO_2_ sensors, and BAG drives omega turns when CO_2_ levels rise^32^. We found that Ca^2+^ responses in URX, BAG and AFD neurons were significantly elevated in *camt-1* mutants across all the O_2_/CO_2_ conditions we tested (Fig. 2a-b, Extended Data Fig. 3a). These data suggest that CAMT-1 activity somehow dampens the Ca^2+^ responses of these sensory neurons. We observed the converse phenotype, dramatically reduced Ca^2+^ levels, when we overexpressed CAMT-1 cDNA specifically in O_2_ sensors or in BAG neurons of control animals (Extended Data Fig. 3b-c). Although Yellow Cameleon is a ratiometric sensor, we cannot exclude the possibility that its reduced expression in animals overexpressing CAMT-1 (Extended Data Fig. 3d) contributes to the reduced baseline YFP/CFP ratio.

**Figure 2:**
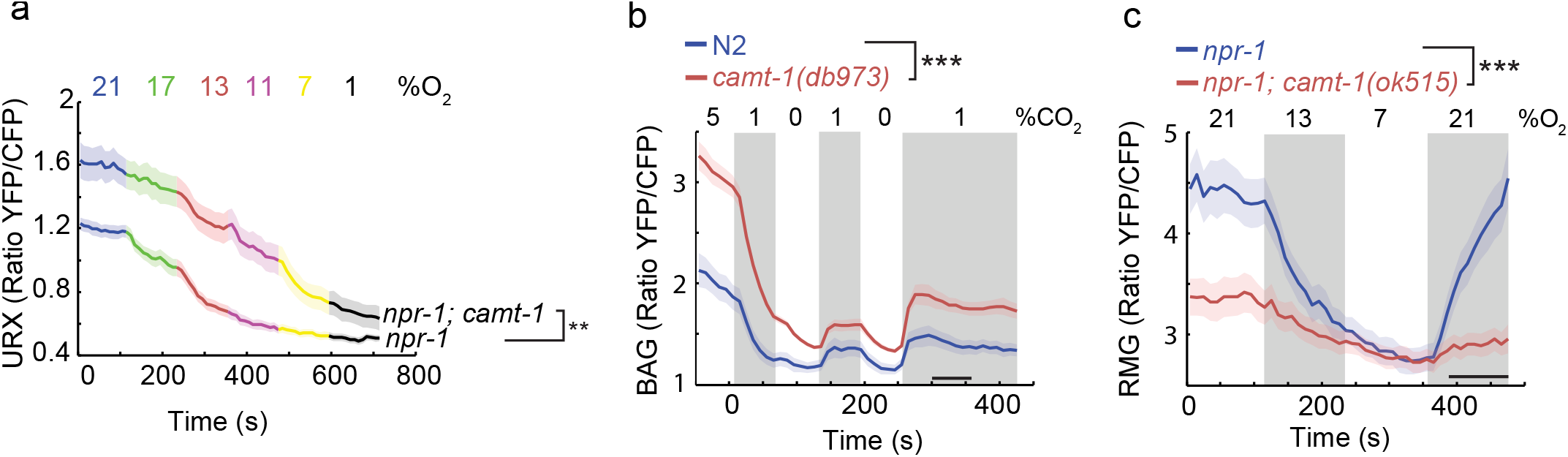
CAMT-1 mutants show heightened stimulus-evoked Ca^2+^ responses in sensory neurons. **a-b** The URX O_2_-sensing neurons (**a**) and the BAG CO_2_ sensors (**b**) show enhanced Ca^2+^ responses across a range of stimulus intensities in *camt-1(db973)* mutants. **c** The amplitude of Ca^2+^ responses to 21% O_2_ are decreased in *camt-1(ok515)* mutants in the interneuron RMG. n ≥ 15 animals (**a**), n ≥ 18 animals (**b**), n ≥ 28 animals (**c**). These strains express a Yellow Cameleon sensor in O_2_-sensing neurons (**a**), BAG (**b**) and RMG (**c**) (see Methods). Average YFP/CFP ratios (line) and SEM (shaded regions) are plotted. *npr-1* and *camt-1* denotes *npr-1(ad609)* and *camt-1(db973)*, respectively. **: p < 0.01, ***: p < 0.001, Mann-Whitney *U* test.

RMG hub interneurons likely integrate inputs from multiple sensory stimuli, including food, pheromones and O_2_, and drive the change in locomotory state associated with a switch from 7% to 21% O_2_^33,34^. O_2_-evoked responses in RMG neurons depend on URX: URX ablation abolishes these responses. Unexpectedly we found that, despite CAMT-1 loss increasing Ca^2+^ levels in URX, Ca^2+^ responses in RMG were strongly reduced in *camt-1* mutants (Fig. 2c). Whether the RMG phenotype reflects a homeostatic response to hyperactivation by URX, or altered input into RMG from other neurons is unclear. However, these data suggest that the reduced locomotory activity of *camt-1* mutants at 21 % O_2_ is due to defective RMG responsiveness.

### CAMT-1 phenotypes are associated with reduced expression of CMD-1/CaM

To identify downstream targets of CAMT-1 we compared the transcriptional profiles of multiple neural types in *camt-1* and control animals^35^. We separately profiled the O_2_-sensors URX/AQR/PQR, the RMG interneurons, the AFD thermosensors, and the BAG O_2_/CO_2_ sensors. We used FACS to collect the neurons from strains in which they were labeled with GFP, and performed 4 – 10 biological replicates for robust statistical power. Our analysis of the data revealed altered expression of many genes; most of these changes were neural-type specific (Fig. 3a, Extended Data Table 1 and 2). A striking exception was the *C. elegans* CaM, *cmd-1. cmd-1* was one of only two genes whose expression was reduced in all four neural profiles relative to WT controls, with mRNA levels 2.5 – 4 fold lower than controls, depending on neural type (Fig. 3a-b). The other gene, Y41C4A.17, is a short transmembrane protein with no known homolog in mammals. CaM plays a key role in regulating many functions in the nervous system^36,37^. We therefore speculated that reduced CMD-1/CaM expression could account for many *camt-1* phenotypes.

**Figure 3:**
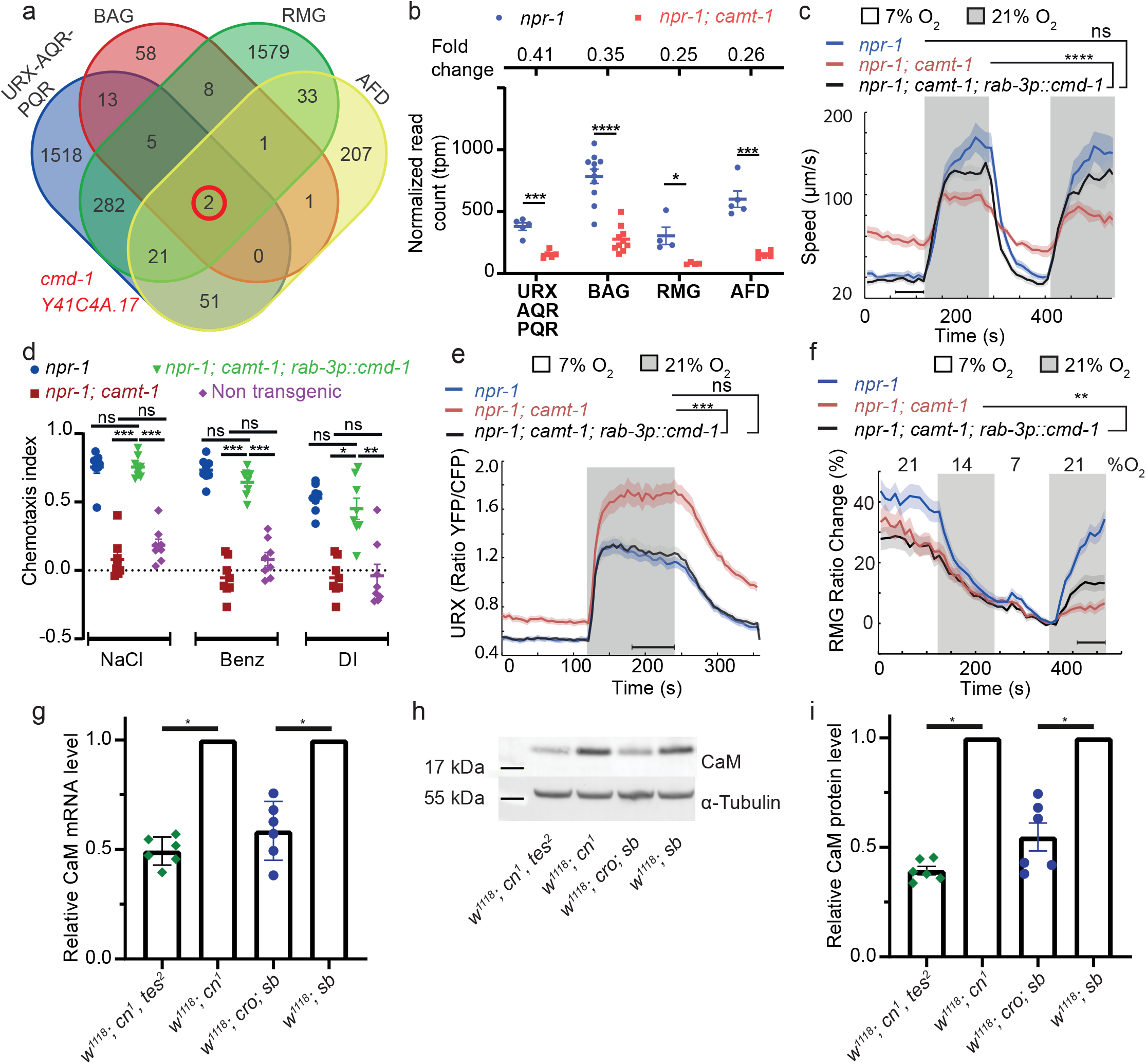
CAMT-1’s pleiotropic phenotypes reflect a conserved role regulating expression of Calmodulin. **a** Venn diagram showing numbers of genes differentially regulated by CAMT-1 in different neuron types we profiled (URX/AQR/PQR, BAG, AFD, RMG). Two genes, *cmd-1 (calmodulin-1)* and *Y41C4A.17*, show consistently altered expression in all neural types profiled. **b** *cmd-1* transcript read counts and fold change (top) for URX/AQR/PQR, BAG, AFD and RMG neurons in *camt-1* mutants compared to controls. Each dot or square represents a separate RNA Seq experiment. **c**-**d** Supplementing CMD-1 expression in neurons using a *rab-3p::cmd-1* transgene rescues the O_2_-response (**c**) and chemotaxis (**d**) phenotypes of *camt-1* mutants. **e**-**f** Supplementing CMD-1 expression in neurons also rescues the O_2_-evoked Ca^2+^-response phenotypes of URX and RMG neurons in *camt-1* mutants. **g** *tes^2^* and *cro* mutants show a decrease in CaM mRNA level compared to control flies. mRNA levels were measured by quantitative PCR. **h-i** *tes^2^* and *cro* mutants show a decrease in CaM protein level compared to control flies. Protein levels were determined using Western blot. (**i**) shows a representative picture and (**i**) shows quantification from multiple samples. ns: p≥0.05, *: p<0.05, **: p < 0.01, ***: p < 0.001, ****: p < 0.0001, Mann-Whitney *U* test (**b-f**), one sample Wilcoxon test to control value of 1 (**g** and **i**). n ≥ 4 replicates for all cell types (**a,b**), n≥ 103 (**c**), n=8 assays for each condition (**d**), n=32 for each genotype (**e**), n≥58 animals (**f**). *npr-1* and *camt-1* denote *npr-1(ad609)* and *camt-1(ok515)* respectively. Lines represent average speed and shaded regions the SEM. For (**c**,**e**-**f**) black horizontal bars indicate time points used for statistical tests

To test this hypothesis, we asked if supplementing CMD-1/CaM expression in *camt-1* mutants, using a pan-neuronal promoter (*rab-3p*), could rescue *camt-1* phenotypes. We made 4 transgenic lines that expressed CMD-1 to different levels (Extended Data Fig. 4a). To monitor expression we placed sequences encoding mCherry in an operon (*SL2*) with *cmd-1*. The *rab-3p::cmd-1::SL2::mCherry* transgene expressing the lowest levels of fluorescence (Line A, Extended Data Fig. 4a) strongly rescued the abnormal O_2_-escape response of *camt-1* mutants (Fig. 3c). Increasing the expression level of CMD-1 restored quiescence behaviour in animals kept at 7% O_2_ but progressively reduced the speed attained at 21% O_2_ (Extended Data Fig. 4b). Supplementing CMD-1 in the nervous system using the lowest expressing line also restored normal chemotaxis towards salt, benzaldehyde and diacetyl in *camt-1* mutants (Fig. 3d). We also found that supplementing CMD-1 levels rescued the hyperexcitability defects in URX and BAG neurons of *camt-1* mutants (Fig. 3e, Extended Data Fig. 4c), and partially rescued Ca^2+^ responses in RMG (Fig. 3f). Together these data suggest that reduced CMD-1 expression accounts for *camt-1* Ca^2+^ signaling and behavioural defects.

### CAMTA promotes CaM expression in *D. melanogaster*

Fly mutants of CAMTA show slow termination of photoresponses compared to wild type controls^38^, and also exhibit defects in male courtship song^26^. An allele of the *Drosophila* calmodulin gene that deletes part of the promoter and reduces CaM expression also shows slow termination of photoresponses^39^. This phenotypic similarity, and our findings in *C. elegans*, prompted us to ask if CAMTA promotes CaM expression in flies too. We obtained two characterized alleles of Drosophila CAMTA (dCamta), *tes^2^* and *cro*, which respectively contain an L1420Stop mutation and a transposon insertion^26,38^. There is a modest decrease in dCAMTA mRNA level in *tes^2^* mutants, suggesting that the premature stop late in the protein may not induce mRNA degradation (Extended Data Fig. 4d). The level of dCAMTA mRNA decreases strongly in *cro* mutants as previously reported^26^ (Extended Data Fig. 4d). We assessed the level of CaM in the heads of dCamta mutant flies using quantitative RT-PCR and Western blots. Each method reported significant decreases in CaM expression in *tes^2^* and *cro* mutants compared to controls (Fig. 3g-i). These results suggest that the transcriptional upregulation of CaM by CAMTA is conserved from worms to flies.

### CAMT-1 directly regulates CMD-1/CaM transcription through multiple binding sites at *cmd-1/CaM* promoter

To test whether CAMT-1 directly binds the *cmd-1* promoter, we performed chromatin immunoprecipitation sequencing (ChIP-seq) using a CRISPR-knockin CAMT-1a::GFP strain. Our analysis revealed > 200 loci that were significantly enriched in CAMT-1a::GFP pulldowns compared to input, and to a mock pulldown (Extended Data Table 3). Importantly, we observed three peaks ~ 6.3kb, 4.8 kb and 2.2 kb upstream of the CMD-1 translation start site in CAMT-1a::GFP pulldown (Fig. 4a, we called these peaks A, B and C respectively, Extended Data Fig. 5a). Thus, CAMT-1 appears to be recruited to multiple sites upstream of *cmd-1*. A CAMT-1 binding peak was also found in the promotor region of Y41C4A.17 (Extended Data Fig. 5b).

To test whether the CAMT-1 ChIP-seq peaks in the *cmd-1* promoter region regulate CMD-1 transcription, we generated CRISPR strains that delete one or more of these peaks. A strain harbouring 110 bp and 136bp deletion at peaks B and C respectively (Fig. 4a-b, *db1275*) and a strain harbouring a 200 bp deletion at peak A (Fig. 4a-b, *db1280*) exhibited aggregation and O_2_ escape responses similar to *npr-1* mutant controls (Fig. 4b). However, a strain harbouring all three deletions (Fig. 4b-c, *db1278*) exhibited strong aggregation defects (Extended Data Fig. 1a) and locomotory responses to O_2_ that mirrored the defects of *camt-1(ok515)* loss-of-function mutants (Fig. 4b, Fig. 1a). Notably, the hyperactivity at 7% O_2_ of *db1278* mutants could be rescued by expressing additional CMD-1 in the nervous system. Like *camt-1(ok515)* mutants, *camt-1(db1278)* mutants also showed chemotaxis defects towards salt, benzaldehyde and diacetyl that could be rescued by neuronal expression of CMD-1 (compare Fig. 3d and 4c). These results suggest that multiple sites in the CMD-1 promoter act redundantly to recruit CAMT-1.

Together our data shows that CAMT-1 regulates CaM expression in a redundant manner through binding to multiple sites in the CaM promoter.

**Figure 4:**
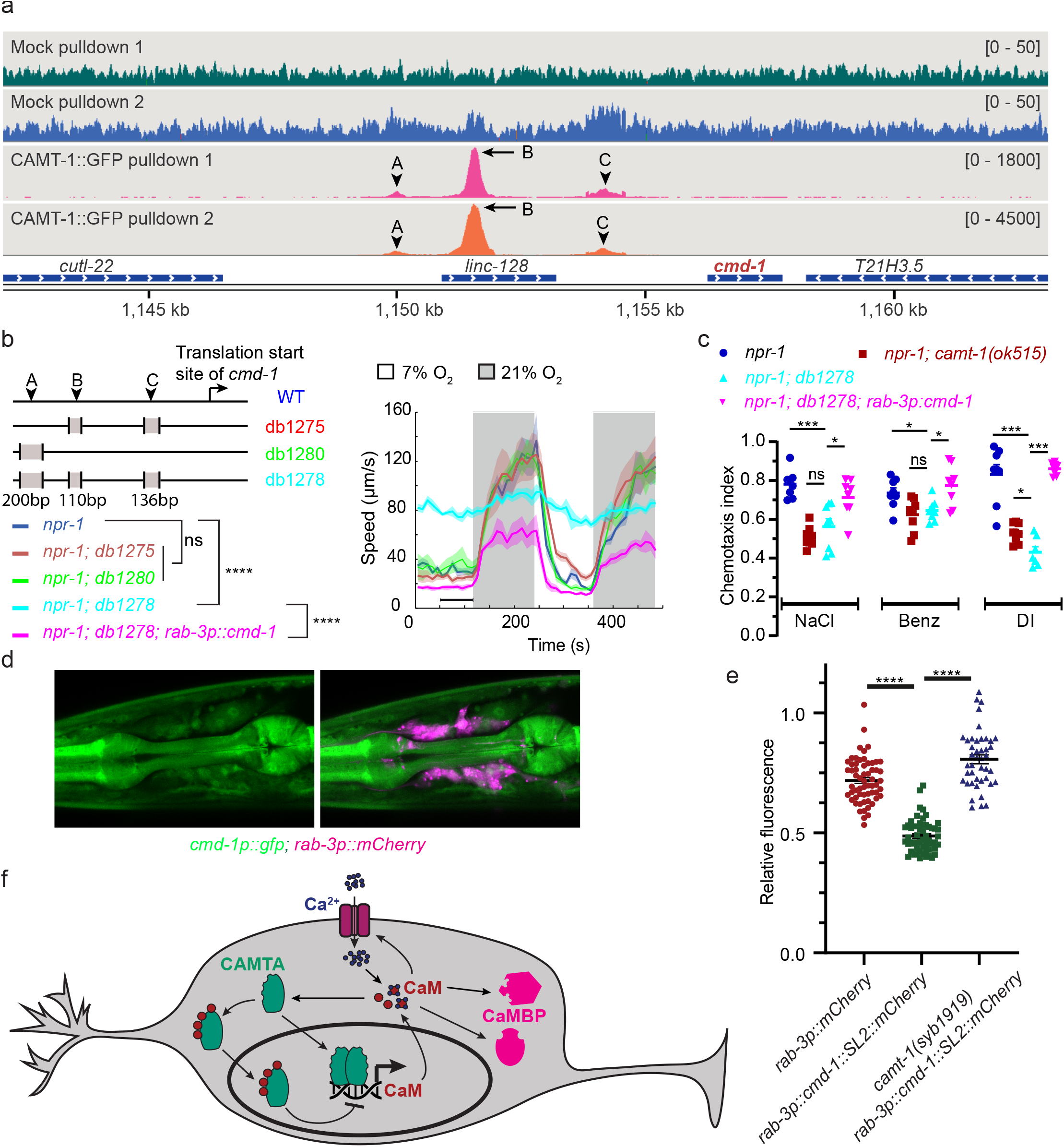
CAMT-1 directly activates CMD-1/CaM expression and can repress CMD-1/CaM expression in the presence of high level of CMD-1/CaM. **a** Coverage plots of chromatin pulldown samples showing enrichment at *cmd-1* promoter in CAMT-1::GFP pulldown (peaks A, B, and C; arrows: major peaks, arrow heads: minor peaks) compared to a mock pulldown or input (Extended Data Figure 5a). Bracketed numbers on the right indicate the scale (normalized read counts). **b** Left, CRISPR-generated strains deleted for one or more of the CAMT-1 ChIP-seq peaks A, B, C shown in **a**; deletions are not drawn to scale. Right, O_2_-evoked speed responses of the promoter deletion strains shown at left. The *db1278* allele in which all three CAMT-1 peaks are deleted confers a strong phenotype that can be rescued by supplementing CMD-1 expression in the nervous system. The *db1275* and *db1280* alleles, which delete only one or two sites have no obvious phenotype. **c** The *db1278* allele confers chemotaxis defects to NaCl, benzaldehyde and diacetyl that can be rescued by supplementing CMD-1 expression in the nervous system. **d** A transcriptional reporter of CMD-1 (*cmd-1p::gfp*) shows expression in neurons, pharyngeal and body wall muscles. Neurons are co-labelled with mCherry driven from *rab-3p*. **e** Over-expression of CMD-1 in neurons (*rab-3p::cmd-1*) decreases the expression of CMD-1 transcriptional reporter (shown in **d**) that can be restored by introducing mutations in the 4 IQ domains of CAMT-1 (the *syb1919* allele). These lines express mCherry in neurons either directly from *rab-3p* or in an operon with CMD-1 (*rab-3p::cmd-1::SL2::mCherry*) and all have *npr-1(ad609)* background mutation. **f** Model of how CAMT-1 positively and negatively regulates levels of CaM in neurons. The binding of four apo-CaM to CAMTA is hypothetical. CaMBP: Other CaM-binding proteins. ns: p≥0.05, *: p<0.05, ***: p < 0.001, ****: p < 0.0001, Mann-Whitney *U* test. n =2 (**a**), n≥ 49 (**b**), n=8 assays for each condition (**c**), n≥ 43 (**e**). *npr-1* denotes *npr-1(ad609)*. Lines represent average speed and shaded regions the SEM. For (**b**) black horizontal bars indicate time points used for statistical tests.

### CMD-1/CaM can inhibit its own expression via CAMT-1

CMD-1 levels are important for proper neural function. We speculated that CMD-1 might homeostatically regulate its own expression by means of a negative feedback loop. To investigate this hypothesis, we built a transcriptional reporter of CMD-1 by fusing an 8.9 kb fragment immediately upstream of the CMD-1 translational start site with GFP. This reporter shows strong fluorescence expression in neurons and muscle, including pharyngeal muscle (Fig. 4d). We crossed this line with another *C. elegans* line that over-expresses CMD-1 in neurons using the *rab-3* promoter (*rab-3p::cmd-1*) and measured neuronal GFP fluorescence in single and double transgenic animals. We normalized expression using pharyngeal GFP levels. Animals bearing both transgenes exhibited a decrease in neuronal GFP fluorescence, suggesting that high levels of CMD-1 can repress expression from the *cmd-1* promoter (Fig. 4e). To examine if this repression is achieved via CaM binding to CAMT-1, we introduced into the double transgenic background the *camt-1* allele that disrupts the 4 IQ domains, *syb1919*. We found that *camt-1(syb1919)* animals expressing *cmd-1p::GFP* and *rab-3p::cmd-1* showed neuronal GFP levels similar to that found in control animals lacking the *rab-3p::cmd-1* transgene (Fig. 4e). These data suggest that CMD-1 can negatively regulate its own expression by binding the IQ domains of CAMT-1. Thus CAMT-1 can not only activate *cmd-1* expression but also repress it when available CMD-1 levels are high.

## Discussion

As CaM is a central regulator of Ca^2+^ signaling, its dysregulation is likely to have profound effects on cellular and organismal processes, and drive of disease^16,36,37^. We discovered that CAMTA is a major regulator of CaM in the nervous system of *C. elegans* and *Drosophila*. The increase in neural excitability and pleiotropic behavioural defects of *camt-1* mutants are consistent with the many known roles of CaM in the nervous system, including controlling activation properties of ion channels^40^ and regulating CaM-dependent kinases ^36,37,41^. Likewise, CaM has been shown to regulate the termination of visual response in *Drosophila^39^*, mirroring the defects of the dCAMTA mutant^38^. Importantly, we uncovered a negative feedback loop to regulate the abundance of CaM via its binding to IQ domains of CAMT-1 (Fig. 4e). Mutant analysis in plants and in flies have implicated CaM/Ca^2+^ binding in the regulation of CAMTA acitivity^3,42,43^. The addition of CaM also decreased transcriptional activity of rice CAMTA in an heterologous system^24^. Our data suggest that binding to CaM converts CAMT-1 from an activator to a repressor, either of its own activity or of that of other transcription activators of CaM. Since CAMTA has several CaM binding sites^1^, it would be interesting to explore whether its activator/repressor activity changes as a function of CaM concentration. Finally, a spectrum of neurological phenotypes, including neurodegeneration and reduced memory performance^14,15^, have been attributed to loss of human CAMTA1. These defects might be attributable reduction in CaM level, given the central role of CaM in the nervous system^36,37^.

## Extended Data Figures

**Extended Data Figure 1. CAMT-1 structure.**

**a** Worm clumps (arrowheads) formed by *npr-1(ad609)* null mutants and disruption of aggregation in *npr-1(ad609); camt-1(ok515)* and *npr-1(ad609); cmd-1p(db1278)* double-mutants.

**b** Sequence alignment of CAMTA proteins, highlighting the conserved CG-1 DNA binding domain (orange), the IPT/TIG Domain (black), Ankyrin repeats (purple), putative Ca^2+^-dependent CaM-binding domain CaMBD (yellow) and IQ region (green). CRISPR/Cas9-generated alleles encoding site-specific point mutations in the CG-1 domain are indicated by orange horizontal bars. CRISPR/Cas9-generated alleles encoding site-specific point mutations of isoleucine in IQ motifs to asparagine are indicated by green horizontal bars. Domain predictions are based on Uniprot and Calmodulin Target Database (http://calcium.uhnres.utoronto.ca/ctdb/ctdb/home.html).

**c** Helical wheel projection of CaM-binding domain (CaMBD) using Emboss peepwheel tool (http://emboss.bioinformatics.nl/cgi-in/emboss/pepwheel?_pref_hide_optional=0) showing a hydrophobic and a positively charged face. Hydrophobic residues are boxed while positively charged residues are marked with a star. Subscript numbers mark two residues R995 and Y996 that are mutated in subsequent rescue experiment (**e**).

**d** *npr-1(ad609); camt-1(syb1919)* mutants with 4 IQ motifs mutated (as shown in **b**) exhibited similar locomotive response to oxygen as *npr-1(ad609)*.

**e** CAMT-1a isoform with mutations R995E and Y996H in the CaMBD (as shown in **c**) still rescue locomotive defects in response to oxygen of *camt-1(ok515)* mutants.

**f-g** Defective responses of *camt-1(ok515)* mutants to 21% O_2_ are rescued by expression of CAMT-1a in RMG (using the *flp-5* promoter, **f**) but not in O_2_-sensing neurons (using the *gcy-32* promoter, **g**). The defective response *camt-1(ok515)* mutants to 7% O_2_ are neither rescued by expression in RMG nor by expression in both RMG and O_2_-sensing neurons (*gcy-32p+flp-5p::camt-1*). # and $ marked the interval used for time points used for statistical test at 7% and 21% O_2_ respectively.

For **d**-**g**, Mann-Whitney *U* test, ns: p≥ 0.05, *: p < 0.05, ****: p < 0.0001, n> 37 (**d**), n ≥ 27 (**e**), n ≥ 28 (**f**), n ≥ 34 animals (**g**), average speed (line) and SEM (shaded regions) are plotted, black horizontal bars show time points used to test.

**Extended Data Figure 2. CAMT-1 is widely expressed in the nervous system.** A CAMT-1a fosmid-based reported, tagged C-terminally with GFP, colocalizes with markers for URX (*gcy-32p::mCherry*, **a**), RMG (*flp-5p::mCherry*, **b**) and ciliated sensory neurons (dil, **c**).

**Extended Data Figure 3**

**a** The AFD neurons, which respond to CO_2_, have enhanced Ca^2+^ levels in *camt-1(db973)* mutants across a range of stimulus intensities.

**b and c** Overexpressing *camt-1* cDNA in O_2_-sensing (using *gcy-32p*, **b**) or BAG neurons (using *flp-17p*, **c**) strongly reduces Ca^2+^ levels.

**d** Quantification of GFP expression driven from the *gcy-37* promoter (same promoter used to drive YC2.60 in Ca^2+^ imaging strain) suggests that YC2.60 expression levels are not significantly affected by disrupting *camt-1*, but are reduced when CAMT-1 is overexpressed.

For **a-c:** These strains express a Yellow Cameleon sensors in AFD (**a**), O_2_-sensing neurons (**b**) and BAG (**c**) (see Methods). Lines represent YFP/CFP ratios and shaded regions the SEM; black bars indicate time points used for statistical tests. ns: p ≥ 0.05, **: p < 0.01, ***: p < 0.001, Mann-Whitney *U* test (**a-c**), ANOVA with Tukey’s post hoc HSD (**d**). n ≥ 15 (**a**), n ≥ 17 (**b**), n ≥ 20 (**c**) and n ≥ 23 animals (**d**).

**Extended Data Figure 4:**

**a-b** Levels of CMD-1 expression, quantified using fluorescence signal from an mCherry expressed in an operon with CMD-1 (*cmd-1::SL2::mCherry*), in different *C. elegans* transgenic lines (**a**), and speed response of these lines to different O_2_ concentrations (**b**). *npr-1* and *npr-1; camt-1* locomotory response were drawn for comparison. *cmd-1::SL2::mCherry* was driven by the *rab-3* neuronal promoter. Note that the speed at 21% oxygen is inversely correlated with the CMD-1 expression level (**a**-**b**). *npr-1* and *camt-1* denote *npr-1(ad609)* and *camt-1(ok515)* respectively.

**c** Pan-neuronal expression of CMD-1 in *camt-1(ok515)* mutants rescues the Ca^2+^ response phenotype in BAG neurons across a range of CO_2_ concentrations. Animals were assayed in 7% O_2_.

**d** Modest and strong decrease in CAMTA mRNA level, determined using quantitative PCR, in *tes^2^* and *cro* mutant flies, respectively, compared to controls.

*:p<0.05, ***:p<0.001, ****:p<0.001, Mann-Whitney *U* test (**a-c**), one sample Wilcoxon test to control value of 1 (**d**).

**Extended Data Figure 5**

**a** Coverage plots for input samples corresponding to the ChIP-seq pulldown samples shown in Fig. 4a. Numbers on the right indicate the scale (normalized read counts).

**b** Coverage plots of pulldown (first four) and input (last four) samples of mock and CAMT-1::GFP showing a peak in the promoter region of the *Y41C4A.17* gene in the CAMT-1::GFP pulldown samples (black arrows). Numbers on the right indicate the scale (normalized read counts).

## Extended Data Tables

**Extended Data Table 1**: The 100 most highly expressed genes (in order of decreasing read counts, in TPM) from neuron-specific RNA profiling.

**Extended Data Table 2**: Genes differentially-expressed in *camt-1* and WT in the profiled neural types.

**Extended Data Table 3**: Genomic locations differentially bound by CAMT-1 identified using the DiffBind algorithm for ChIP-seq data with a False Discovery Rate (FDR) threshold of 0.05. Genes overlapping or within 10kb downstream of these sites are reported. The table is sorted in the order of increasing FDR. Note that the CAMT-1 binding site at the *cmd-1* promoter was annotated with the overlapping long intervening non-coding RNA *linc-128*.

**Extended Data Table 4**: List of *C. elegans* strains used in this study.

## Methods

No statistical methods were used to predetermine sample size. The experiments were not randomized.

### Strains

*C. elegans* strains used are listed in Extended Data Table 4. Strains were maintained at room temperature, on nematode growth medium (NGM) with *E. coli* OP50. RB746 *camt-1(ok515)*, and OH10689 *otIs355[rab-3p::2xNLS::TagRFP]* were obtained from the *Caenorhabditis* Genetic Center (P40 OD010440).

### Molecular Biology

We obtained a clone containing the *camt-1* locus from the *C. elegans* fosmid library (Source BioScience). To insert GFP immediately prior to the termination codon of *camt-1* we followed established protocols^32^. The primers used to amplify the recombineering cassette from pBALU1 were: ATCATCCATGGGACCAATTGAAACCGCCGTATGGTTGCGGAACACTTGCAATGAGTAAAGGAGAAGAACTTTTCAC and aaaccaataaaaaaaatcggcatcttctaaaagtgacaccggggcaaTTATTTGTATAGTTCATCCATGCCATG. To generate transgenic lines, we injected a mix of 50 ng/μl fosmid DNA and 50 ng/μl co-injection marker (unc-122p::dsRED).

*C. elegans* expression constructs were generated using MultiSite Gateway Recombination (Invitrogen) or FastCloning^44^. We amplified cDNA corresponding to *camt-1 (T05C1.4b)* using primers ggggACAAGTTTGTACAAAAAAGCAGGCTtttcagaaaaATGAATAATTCAGTCACTCGTCTTCTTTTCAAACGACTGCTGAC and ggggACCACTTTGTACAAGAAAGCTGGGTATTATGCAAGTGTTCCGCAACCAT ACGGCG. We were unable to amplify *camt-1* cDNA corresponding to the longer *T05C1.4a* splice variant so we generated it by site-directed mutagenesis of *T05C1.4b* cDNA. To convert *T05C1.4b* cDNA to *T05C1.4a* we used the Q5 Site-Directed Mutagenesis Kit (NEB) and primers gtcatactcaacatctaATTGCGGAAAATGCATGC and catcatcaatatttacaTTATTACGATTTTGTCGCATAAAATTC

### Genome editing

Strains PHX994 and PHX1919 were generated by SunyBiotech on our request (Fu Jian, China). We generated point mutations in the endogenous *camt-1* locus using published CRISPR protocols^45^. Cas9 endonuclease, crRNA and tracrRNA were obtained from IDT (Iowa, US).

### Behaviour assays

O_2_- and CO_2_-response assays were performed as described previously^46^, using young adults raised at room temperature. 15-30 young adults were assayed in a microfluidic PDMS chamber on an NGM plate seeded with 20-50 μl OP50. Indicated O_2_/CO_2_ mixtures (in nitrogen) were bubbled through H2O and pumped into the PDMS chamber using a PHD 2000 Infusion syringe pump (Harvard Apparatus). Videos were recorded at 2 fps using FlyCapture software (FLIR Systems), and a Point Gray Grasshopper camera mounted on a Leica MZ6 microscope. Custom MATLAB software (Zentracker: https://github.com/wormtracker/zentracker) was used to measure speed and omega turns.

Chemotaxis assays were performed as previously described (Bargmann et al., 1993) with minor modifications. 9 cm assay plates were made with 2% Bacto Agar, 1mM CaCl_2_, 1mM MgSO_4_ and 25mM K_2_HPO_4_ pH 6. Test and control circles of 3cm diameter were marked on opposite sides of the assay plate, equidistant from a starting point where >50 animals were placed to begin the assay. For olfactory assays, 1μl odorant (Benzaldehyde 1/400 or Diacetyl 1/1000 dilution in ethanol) or 1μl ethanol, and 1μl 1M NaN3, was added to each circle.

For gustatory assays, an agar plug containing 100 mM NaCl was added the night before and removed prior to assay. Assays were allowed to proceed for 30-60min, after which point plates were moved to 4°C. Chemotaxis index was calculated as (number of animals in test circle – number of animals in control circle) / total number of animals that have left starting area.

### Heat-shock

Animals were raised at 15 °C to reduce leaky expression from the *hsp-16.41* heat-shock promoter. To induce heat-shock, parafilm-wrapped plates were submerged in a 34 °C water bath for 30 min, and then recovered at room temperature for 6 h.

### Ca^2+^ imaging

Neural imaging was performed as previously described^46^, with a ×2 AZ-Plan Fluor objective (Nikon) on a Nikon AZ100 microscope fitted with ORCA-Flash4.0 digital cameras (Hamamatsu). Excitation light was provided from an Intensilight C-HGFI (Nikon), through a 438/24 nm filter and an FF458DiO2 dichroic (Semrock). Emission light was split using a TwinCam dual camera adapter (Cairn Research) bearing a filtercube containing a DC/T510LPXRXTUf2 dichroic and CFP (483/32 nm) and YFP (542/27) filters. We acquired movies using NIS-Elements (Nikon), with 100 or 500 ms exposure time. YFP/CFP ratios in URX were reported by YC2.60 driven from the *gcy-37* promoter, in BAG by YC3.60 driven from the *flp-17* promoter, in AFD by YC3.60 driven from the *gcy-8* promoter, and in RMG by a strain expressing *ncs-1p::cre* and *pflp-21::LoxP-STOP-LoxP::YC2.60*.

### Single-neuronal-type cell sorting and RNA sequencing

We used *C. elegans* lines in which neuronal types were labelled by expressing GFP under specific promoters: oxygen sensing neurons (*gcy-37p*), BAG (*flp-17p*), RMG (combination of *ncs-1p::CRE* and *flp-21::loxP::STOP::loxP::GFP^34^*) and AFD (*gcy-8p*). These markers were crossed into either *npr-1(ad609)* or *npr-1(ad609); camt-1(ok515)* backgrounds. *C. elegans* cells were dissociated and GFP-labelled cells sorted as described previously^35^. Briefly, *C. elegans* with GFP-labelled neurons were synchronized using the standard bleaching protocol and eggs placed 3 days before cell sorting on 90 mm NGM plates seeded with OP50. For each sample, we used >50 000 worms. The worms were washed 3 times with M9, prewashed and then incubated for 6.5 min with 750 μl lysis buffer (0.25% SDS, 200 mM DTT, 20 mM HEPES pH 8.0, 3% sucrose). The worms were then rapidly washed 5 times with M9. We dissociated the cells by adding 500 μl of Pronase (Roche) 20 mg/ml and by either pipetting up-and-down or stirring continuously for 12 min using a small magnetic stirrer. The pronase were inactivated by adding 500 μl of PBS + 2% Fetal Bovine Serum (GIBCO). The solutions were passed through a 5 μm pore size syringe filter (Millipore), and filtered cells further diluted in PBS + 2% FBS for sorting using a Sony Biotechnology Synergy High Speed Cell Sorter. Gates for detection were determined using cells prepared in parallel from non-fluorescent animals using the same protocol. An average of 3000 cells were collected for each library, and sorted directly into lysis buffer containing RNAse inhibitor (NEB E6420). cDNA libraries were made from RNA using NEB’s Next Single Cell/Low Input RNA Library Prep Kit for Illumina (NEB E6420). Libraries were sequenced on an Illumina HiSeq 4000 with single reads of 50 bases.

### Confocal microscopy and image analysis

Young adult worms were mounted for microscopy on a 2% agar pad in 1M sodium azide. Image analysis and fluorescence quantification was carried out using Fiji (ImageJ, Wayne Rasband, NIH).

The expression pattern of CAMT-1(fosmid)-GFP was imaged as previously described^46^ on an Inverted Leica SP8 confocal microscope using a 63x/1.20 N.A. water-immersion objective. Colocalization of CAMT-1(fosmid)-GFP with neuronal markers was imaged on a Andor Ixon EMCCD camera coupled to a spinning disk confocal unit, with a 60x lens and 100 ms exposure time..

Lines expressing a *cmd-1* transcriptional reporter (*cmd-1p::gfp*) and a red neuronal marker (either *rab-3p::mCherry* or *rab-3p::cmd-1::SL2::mCherry*) were imaged on an LSM800 inverted microscope (Zeiss) using a 63x/1.40 N.A. oil-immersion objective. The region between the two pharyngeal bulbs (Fig. 4a) were imaged using stacks of 0.3 μm in step size. A section of 3 μm (10 images) around the middle of the pharynx was projected using the maximum projection method. The neurons were identified by thresholding the intensity of the red marker (mCherry). The neuronal region overlapping with the pharynx or body wall muscles is excluded. The relative fluorescence in Fig. 4e was defined as the GFP level in neurons minus background fluorescence divided by the level of fluorescence in the pharynx (metacorpus + isthmus + terminal bulb) minus background fluorescence.

### ChIP-seq

The ChIP-seq protocol used was similar to that describe in Wormbook (http://www.wormbook.org/chapters/www_chromatinanalysis/chromatinanalysis.html). Briefly, mixed-stage worms were grown in liquid culture, harvested, washed 3 times in PBS and resuspended in PBS + Protease Inhibitor (PI, Sigma). Worm ‘popcorn’ was prepared by dripping worm solution into liquid nitrogen, then hand ground to a fine powder. For each ChIP replicate we used 2.5 g of packed worms. Cross-linking was carried out by incubating samples in 1.5 mM EGS in PBS for 10 min, then adding 1.1% formaldehyde and incubating for a further 10 min. The reaction was quenched using 0.125 M glycine. The pellet was washed once in PBS + PMSF 1mM and once in FA buffer (50 mM HEPES/KOH pH 7.5, 1 mM EDTA, 1% Triton™ X-100, 0.1% sodium deoxycholate, 150 mM NaCl) + PI. The pellet was resuspended in 4ml of FA buffer + PI + 0.1% sarkosyl and sonicated using a Diagenode Bioruptor Plus with 40 cycles, 30 s on, 30 s off. The sample was then spun in a table top microcentrifuge at top speed (15000 rpm) for 15 min. The supernatant was incubated with 1 μl of anti-GFP antibody (Abcam ab290) overnight at 4 °C. 60 μl of Protein A conjugated Dynabeads was added and the resulting solution incubated for 3 h at 4 °C. Pulldown, washing and de-crosslinking steps were as described in http://www.wormbook.org/chapters/www_chromatinanalysis/chromatinanalysis.html. For preparing ChIP libraries, we use NEBNext Ultra II DNA Library Prep Kit for Illumina with half of the pulldown and 30 ng of input. DNA libraries were then sequenced on a Illumina HiSeq 4000 platform with single read of 50 bases.

### RNA-seq and ChIP-seq data analysis

RNA-seq data were mapped using PRAGUI - a Python 3-based pipeline for RNA-seq data analysis available at https://github.com/lmb-seq/PRAGUI. PRAGUI integrates RNA-seq processing packages including Trim Galore, FastQC, STAR, DESeq2, HTSeq, Cufflinks and MultiQC. Output from PRAGUI was analyzed using PEAT - Pragui Exploratory Analysis Tool (https://github.com/lmb-seq/PEAT) to obtain the list of differently expressed genes with a false discovery rate < 0.05. The Venn diagram was drawn using the online tool http://bioinformatics.psb.ugent.be/webtools/Venn/.

ChIP-seq data were analyzed using a nucleome processing and analysis toolkit which contains an automated ChIP-seq processing pipeline using Bowtie2 mapping and MACS2 peak calling. The software is available on Github at https://github.com/tjs23/nuc_tools. Comparisons between different ChIP-seq conditions were carried out using the DiffBind package^47^. ChIP-seq processed data was visualized using IGV^48,49^.

### Fly genetics

*(w^1188^), (w^1118^; cn^1^, tes^2^/cyo)* and *(w^1118^; cro/cyo; sb/TM3 ser)* flies were generously obtained from Daria Siekhaus (IST Austria), Hong-Sheng Li (UMass) and Daisuke Yamamoto (NICT) respectively. *cn^1^* flies was obtained from the Bloomington *Drosophila* Stock Center (NIH P40OD018537). These flies were crossed to obtain *w^1118^; cn^1^* and *w^1118^; sb* control flies.

### Quantitative PCR

qPCR was performed using the Janus Liquid Handler (PerkinElmer) and a LightCycler 480 system (Roche). Total RNA was extracted from the heads of 20 male adults or 17 female adults using a Monarch Total RNA Miniprep Kit (NEB). 3 replicates for male and 3 replicates for female flies were done for each genotype. RNA was reverse transcribed into cDNA using a ImProm-II Reverse Transcription System (Promega). cDNA was mixed with Luna Universal qPCR Master Mix (NEB). *RpL32 (rp49)* was amplified as an internal control. Primer sequences for *Rpl32* and *CAMTA* were identical with the one used in Sato *et al.^26^. CaM* was amplified using the primer pair 5’-TGCAGGACATGATCAACGAG-3’ (forward) and 5’-ATCGGTGTCCTTCATTTTGC-3’ (reverse). Data processing was performed using LightCycler Software (Roche).

### Western Blot

Protein from the heads of ~50 female and 60 male adult flies were extracted using RIPA buffer (150 mM NaCl, 1% NP40, 0.5% sodium deoxycholate, 0.1% SDS, 50 mM Tris-HCl, pH 8.0, Protease inhibitors). 3 replicates for male and 3 replicates for female flies were performed for each genotype. After SDS-PAGE using Bolt 4–12% Bis-Tris Plus gels (ThermoFisher Scientific), protein was transferred to PVDF membrane (0.45-micron pore size, ThermoFisher Scientific) using the TE 22 Mighty Small Transfer Unit (Amersham Biosciences). Membranes were blocked with casein blocking buffer (1% Hammersten Casein, 20mM Tris-HCl, 137 mM NaCl) for 1 h, then incubated with primary antibody overnight at 4 °C, followed by secondary antibody for 1 h at RT. Unbound antibody was washed away with TBS-T or TBS (3 ×5 min). α-tubulin was used as an internal control. The following commercially available antibodies were used: anti-CaM (Abcam, ab45689, diluted 1/500), anti-α-tubulin (Abcam, ab40742, diluted 1/5000), goat anti-rabbit StarBright Blue 700 (Biorad, 12004161, diluted 1/5000) and goat anti-mouse StarBright Blue 520 (Biorad, 12005867, diluted 1/5000). Blots were imaged using Chemidoc MP Imaging System (Biorad).

### Statistical tests

Statistical tests are two-tailed and were performed using Matlab (MathWorks, MA, US), GraphPad Prism (GraphPad Software, CA, US) or R (R Foundation for Statistical Computing, Vienna, Austria, http://www.R-project.org/). Measurements were done from distinct samples.

## Acknowledgement

We thank the MRC-LMB Flow Cytometry facility and Imaging Service for support, the Cancer Research UK Cambridge Institute Genomics Core for Next Generation Sequencing, Julie Ahringer and Alex Appert for advice and technical help for ChIP-seq experiments, Paula Freire-Pritchett, Tim Stevens and Gurpreet Ghattaoraya for RNA-seq and ChIP-seq analysis, Hong-Sheng Li and Daisuke Yamamoto for generously sending the *tes^2^* and *cro* mutants, Daria Siekhaus for hosting the fly work, Michaela Misova for technical assistance. This work was supported by an Advanced ERC grant (269058 ACMO) and a Wellcome Investigator Award (209504/Z/17/Z) to M.d.B, and an IST Plus Fellowship to T.V-B (Marie Sklodowska-Curie agreement No 754411).

## Contributions

T.V-B, S.F and M.d.B. designed experiments. T.V-B and S.F performed experiments and analysed the data. T.V-B, S.F and M.d.B. wrote the paper.

## Corresponding author

Correspondence to Mario de Bono.

## Ethics declarations Competing interests

The authors declare no competing financial interests.

## Supplementary information

This file contains 5 Extended Data Figures and 4 Extended Data Tables.

## Data availability

The datasets generated during current study are either included in this article or deposited at www.geo. (ChIP-seq data) or available from the corresponding author on reasonable request.

## Notes

### Competing Interest Statement

The authors have declared no competing interest.

## References

1 Finkler, A., Ashery-Padan, R. & Fromm, H. CAMTAs: calmodulin-binding transcription activators from plants to human. FEBS Lett 581, 3893–3898, doi:10.1016/j.febslet.2007.07.051 (2007).

2 Yang, T. & Poovaiah, B. W. A calmodulin-binding/CGCG box DNA-binding protein family involved in multiple signaling pathways in plants. J Biol Chem 277, 45049–45058, doi:10.1074/jbc.M207941200 (2002).

3 Du, L. et al. Ca(2+)/calmodulin regulates salicylic-acid-mediated plant immunity. Nature 457, 1154–1158, doi:10.1038/nature07612 (2009).

4 Doherty, C. J., Van Buskirk, H. A., Myers, S. J. & Thomashow, M. F. Roles for Arabidopsis CAMTA transcription factors in cold-regulated gene expression and freezing tolerance. Plant Cell 21, 972–984, doi:10.1105/tpc.108.063958 (2009).

5 Pandey, N. et al. CAMTA 1 regulates drought responses in Arabidopsis thaliana. BMC Genomics 14, 216, doi:10.1186/1471-2164-14-216 (2013).

6 Shkolnik, D., Finkler, A., Pasmanik-Chor, M. & Fromm, H. CALMODULIN-BINDING TRANSCRIPTION ACTIVATOR 6: A Key Regulator of Na(+) Homeostasis during Germination. Plant Physiol 180, 1101–1118, doi:10.1104/pp.19.00119 (2019).

7 Song, K. et al. The transcriptional coactivator CAMTA2 stimulates cardiac growth by opposing class II histone deacetylases. Cell 125, 453–466, doi:10.1016/j.cell.2006.02.048 (2006).

8 Long, C. et al. Ataxia and Purkinje cell degeneration in mice lacking the CAMTA1 transcription factor. Proc Natl Acad Sci U S A 111, 11521–11526, doi:10.1073/pnas.1411251111 (2014).

9 Bas-Orth, C., Tan, Y. W., Oliveira, A. M., Bengtson, C. P. & Bading, H. The calmodulin-binding transcription activator CAMTA1 is required for longterm memory formation in mice. Learning & memory (Cold Spring Harbor, N.Y.) 23, 313–321, doi:10.1101/lm.041111.115 (2016).

10 Huentelman, M. J. et al. Calmodulin-binding transcription activator 1 (CAMTA1) alleles predispose human episodic memory performance. Hum Mol Genet 16, 1469–1477, doi:10.1093/hmg/ddm097 (2007).

11 Thevenon, J. et al. Intragenic CAMTA1 rearrangements cause non-progressive congenital ataxia with or without intellectual disability. J Med Genet 49, 400-408, doi:10.1136/jmedgenet-2012-100856 (2012).

12 Shinawi, M., Coorg, R., Shimony, J. S., Grange, D. K. & Al-Kateb, H. Intragenic CAMTA1 deletions are associated with a spectrum of neurobehavioral phenotypes. Clin Genet 87, 478–482, doi:10.1111/cge.12407 (2015).

13 Faas, G. C., Raghavachari, S., Lisman, J. E. & Mody, I. Calmodulin as a direct detector of Ca2+ signals. Nat Neurosci 14, 301–304, doi:10.1038/nn.2746 (2011).

14 Baimbridge, K. G., Celio, M. R. & Rogers, J. H. Calcium-binding proteins in the nervous system. Trends Neurosci 15, 303–308, doi:10.1016/0166-2236(92)90081-i (1992).

15 Hoeflich, K. P. & Ikura, M. Calmodulin in action: diversity in target recognition and activation mechanisms. Cell 108, 739–742, doi:10.1016/s0092-8674(02)00682-7 (2002).

16 Berchtold, M. W. & Villalobo, A. The many faces of calmodulin in cell proliferation, programmed cell death, autophagy, and cancer. Biochim Biophys Acta 1843, 398–435, doi:10.1016/j.bbamcr.2013.10.021 (2014).

17 Luby-Phelps, K., Hori, M., Phelps, J. M. & Won, D. Ca(2+)-regulated dynamic compartmentalization of calmodulin in living smooth muscle cells. J Biol Chem 270, 21532–21538, doi:10.1074/jbc.270.37.21532 (1995).

18 Pepke, S., Kinzer-Ursem, T., Mihalas, S. & Kennedy, M. B. A dynamic model of interactions of Ca2+, calmodulin, and catalytic subunits of Ca2+/calmodulin-dependent protein kinase II. PLoS Comput Biol 6, e1000675, doi:10.1371/journal.pcbi.1000675 (2010).

19 Chen, C. et al. IL-17 is a neuromodulator of Caenorhabditis elegans sensory responses. Nature 542, 43–48, doi:10.1073/pnas.1618934114 (2017).

20 Busch, K. E. et al. Tonic signaling from O(2) sensors sets neural circuit activity and behavioral state. Nat Neurosci 15, 581–591, doi:10.1371/journal.pgen.1004011 (2012).

21 Rogers, C., Persson, A., Cheung, B. & de Bono, M. Behavioral motifs and neural pathways coordinating O2 responses and aggregation in C. elegans. Current biology: CB 16, 649–659, doi:10.1016/j.cub.2006.03.023 (2006).

22 de Bono, M., Bargmann, C. I., de Bono, M. & Hodgkin, J. Natural variation in a neuropeptide Y receptor homolog modifies social behavior and food response in C. elegans Evolution of sex determination in caenorhabditis: unusually high divergence of tra-1 and its functional consequences. Cell 94, 679–689 (1998).

23 Bouche, N., Scharlat, A., Snedden, W., Bouchez, D. & Fromm, H. A novel family of calmodulin-binding transcription activators in multicellular organisms. J Biol Chem 277, 21851–21861, doi:10.1074/jbc.M200268200 (2002).

24 Choi, M. S. et al. Isolation of a calmodulin-binding transcription factor from rice (Oryza sativa L.). J Biol Chem 280, 40820–40831, doi:10.1074/jbc.M504616200 (2005).

25 Consortium, C. e. D. M. large-scale screening for targeted knockouts in the Caenorhabditis elegans genome. G3 (Bethesda) 2, 1415–1425, doi:10.1534/g3.112.003830 (2012).

26 Sato, K., Ahsan, M. T., Ote, M., Koganezawa, M. & Yamamoto, D. Calmodulin-binding transcription factor shapes the male courtship song in Drosophila. PLoS Genet 15, e1008309, doi:10.1371/journal.pgen.1008309 (2019).

27 Bretscher, A. J., Busch, K. E. & de Bono, M. A carbon dioxide avoidance behavior is integrated with responses to ambient oxygen and food in Caenorhabditis elegans. Proc Natl Acad Sci U S A 105, 8044–8049 (2008).

28 Ward, S. Chemotaxis by the nematode Caenorhabditis elegans: identification of attractants and analysis of the response by use of mutants. Proc Natl Acad Sci US A 70, 817–821 (1973).

29 Bargmann, C. I., Hartwieg, E. & Horvitz, H. R. Odorant-selective genes and neurons mediate olfaction in C. elegans. Cell 74, 515–527, doi:10.1016/0092-8674(93)80053-h (1993).

30 West, A. E., Griffith, E. C. & Greenberg, M. E. Regulation of transcription factors by neuronal activity. Nature reviews. Neuroscience 3, 921–931, doi:10.1038/nrn987 (2002).

31 Chin, D. & Means, A. R. Calmodulin: a prototypical calcium sensor. Trends Cell Biol 10, 322–328, doi:10.1016/s0962-8924(00)01800-6 (2000).

32 Bretscher, A. J. et al. Temperature, oxygen, and salt-sensing neurons in C. elegans are carbon dioxide sensors that control avoidance behavior. Neuron 69, 1099–1113, doi:10.1073/pnas.1106134109 (2011).

33 Laurent, P. et al. Decoding a neural circuit controlling global animal state in C. elegans. Elife 4 (2015).

34 Macosko, E. Z. et al. A hub-and-spoke circuit drives pheromone attraction and social behaviour in C. elegans. Nature 458, 1171–1175, doi:10.1038/nature07886 (2009).

35 Kaletsky, R. et al. Transcriptome analysis of adult Caenorhabditis elegans cells reveals tissue-specific gene and isoform expression. PLoS Genet 14, e1007559, doi:10.1371/journal.pgen.1007559 (2018).

36 Wayman, G. A., Lee, Y. S., Tokumitsu, H., Silva, A. J. & Soderling, T. R. Calmodulin-kinases: modulators of neuronal development and plasticity. Neuron 59, 914–931 (2008).

37 Zalcman, G., Federman, N. & Romano, A. CaMKII Isoforms in Learning and Memory: Localization and Function. Front Mol Neurosci 11, 445 (2018).

38 Han, J. et al. The fly CAMTA transcription factor potentiates deactivation of rhodopsin, a G protein-coupled light receptor. Cell 127, 847–858, doi:10.1016/j.cell.2006.09.030 (2006).

39 Scott, K., Sun, Y., Beckingham, K. & Zuker, C. S. Calmodulin regulation of Drosophila light-activated channels and receptor function mediates termination of the light response in vivo. Cell 91, 375–383, doi:10.1016/s0092-8674(00)80421-3 (1997).

40 Saimi, Y. & Kung, C. Calmodulin as an ion channel subunit. Annual review of physiology 64, 289–311, doi:10.1146/annurev.physiol.64.100301.111649 (2002).

41 Xia, Z. & Storm, D. R. The role of calmodulin as a signal integrator for synaptic plasticity. Nature reviews. Neuroscience 6, 267–276, doi:10.1038/nrn1647 (2005).

42 Nie, H. et al. SR1, a calmodulin-binding transcription factor, modulates plant defense and ethylene-induced senescence by directly regulating NDR1 and EIN3. Plant Physiol 158, 1847–1859 (2012).

43 Kim, Y. S. et al. CAMTA-Mediated Regulation of Salicylic Acid Immunity Pathway Genes in Arabidopsis Exposed to Low Temperature and Pathogen Infection. Plant Cell 29, 2465–2477 (2017).

44 Li, C. et al. FastCloning: a highly simplified, purification-free, sequence- and ligation-independent PCR cloning method. BMC Biotechnol 11, 92, doi:10.1186/1472-6750-11-92 (2011).

45 Dokshin, G. A., Ghanta, K. S., Piscopo, K. M. & Mello, C. C. Robust Genome Editing with Short Single-Stranded and Long, Partially Single-Stranded DNA Donors in Caenorhabditis elegans. Genetics 210, 781–787 (2018).

46 Flynn, S. M. et al. MALT-1 mediates IL-17 neural signaling to regulate C. elegans behavior, immunity and longevity. Nat Commun 11, 2099, doi:10.1038/s41467-020-15872-y (2020).

47 Stark, R. & Brown, G. DiffBind: differential binding analysis of ChIP-Seq peak data. Bioconductor (2011).

48 Robinson, J. T. et al. Integrative genomics viewer. Nat Biotechnol 29, 24–26, doi:10.1038/nbt.1754 (2011).

49 Thorvaldsdottir, H., Robinson, J. T. & Mesirov, J. P. Integrative Genomics Viewer (IGV): high-performance genomics data visualization and exploration. Brief Bioinform 14, 178–192, doi:10.1093/bib/bbs017 (2013).

