## Extended Data Figures for "Transcriptional Control Of Calmodulin By CAMTA Regulates Neural Excitability"

Extended Data Figure 1

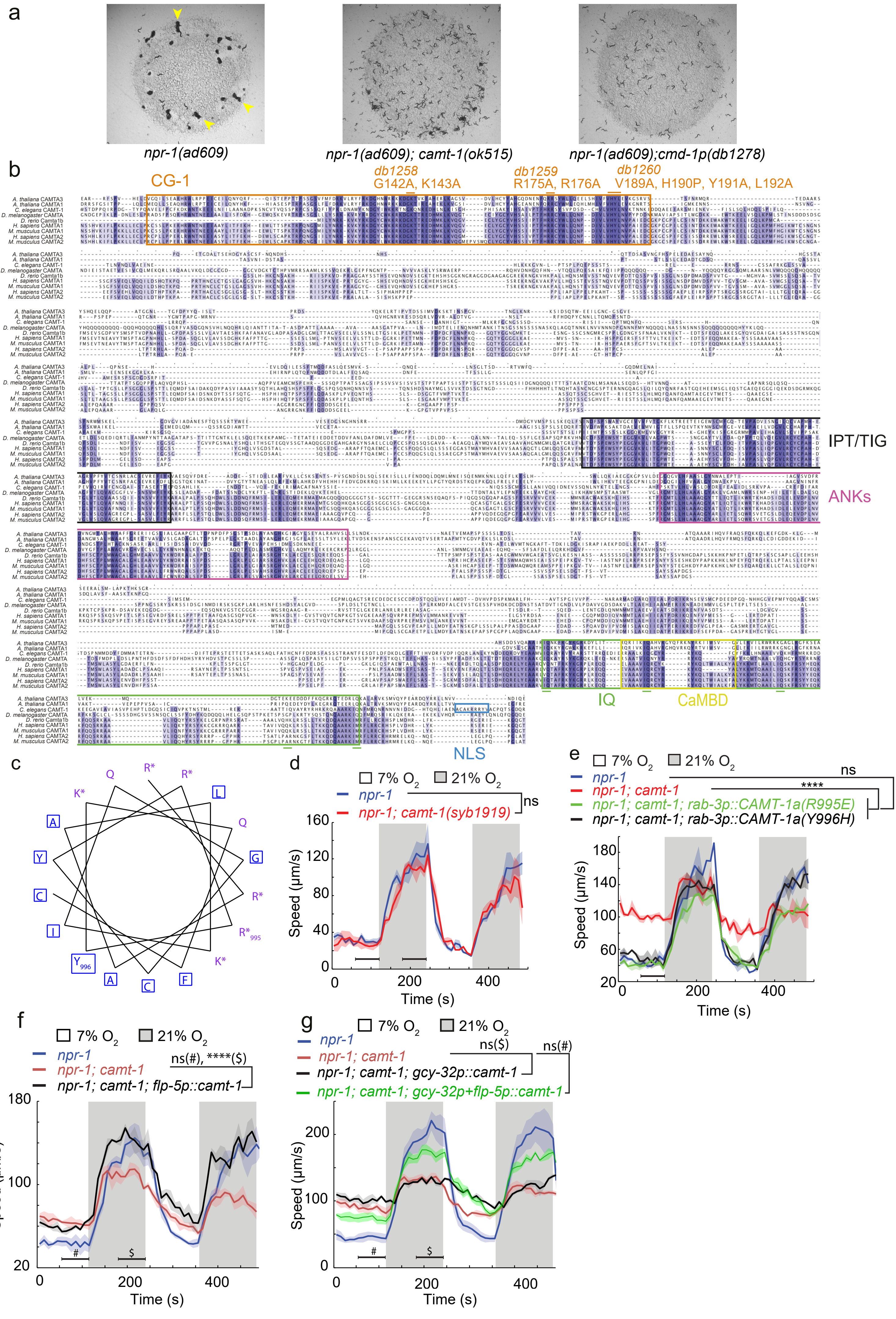

Extended Data Figure 2

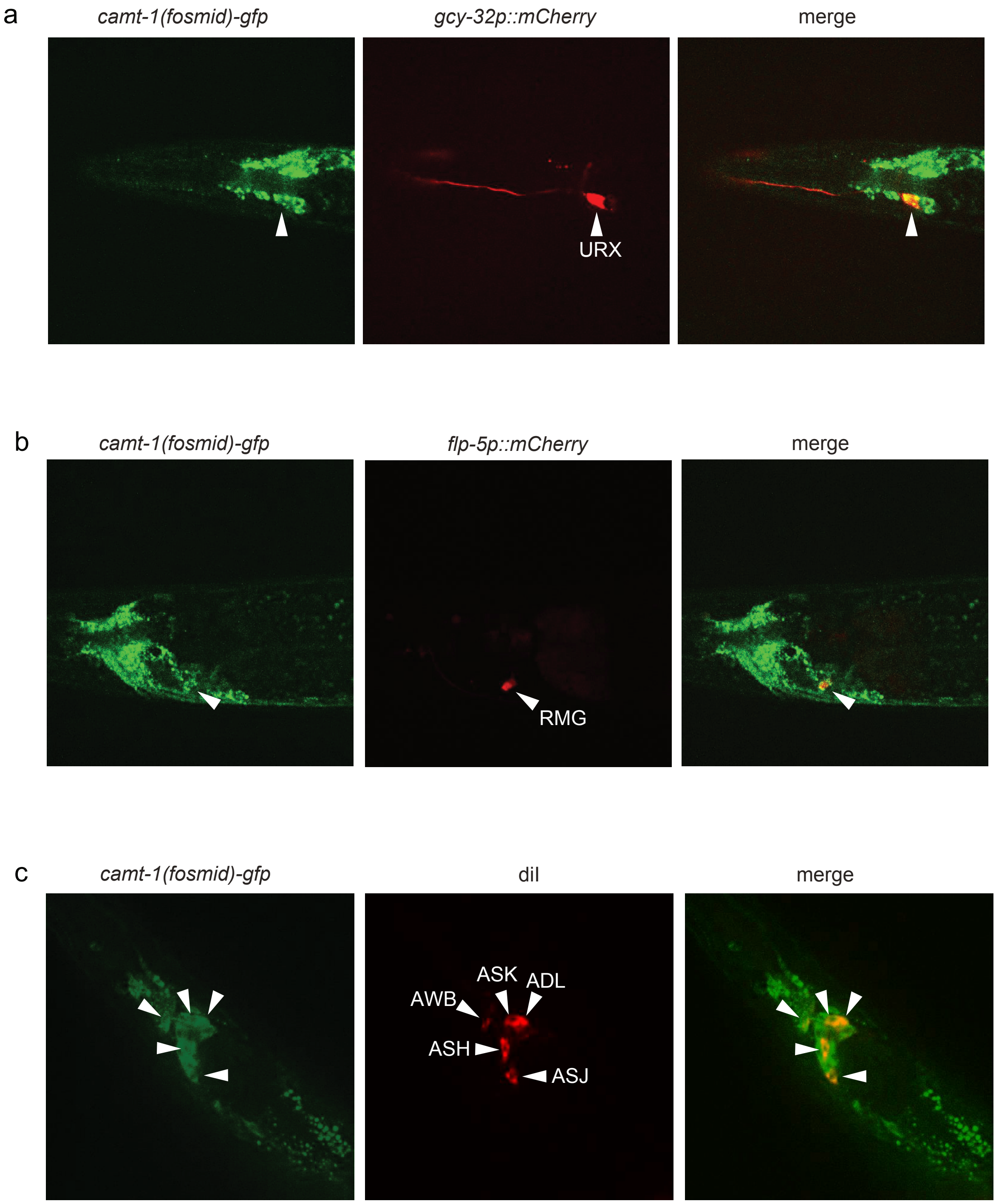

### Extended Data Figure 3

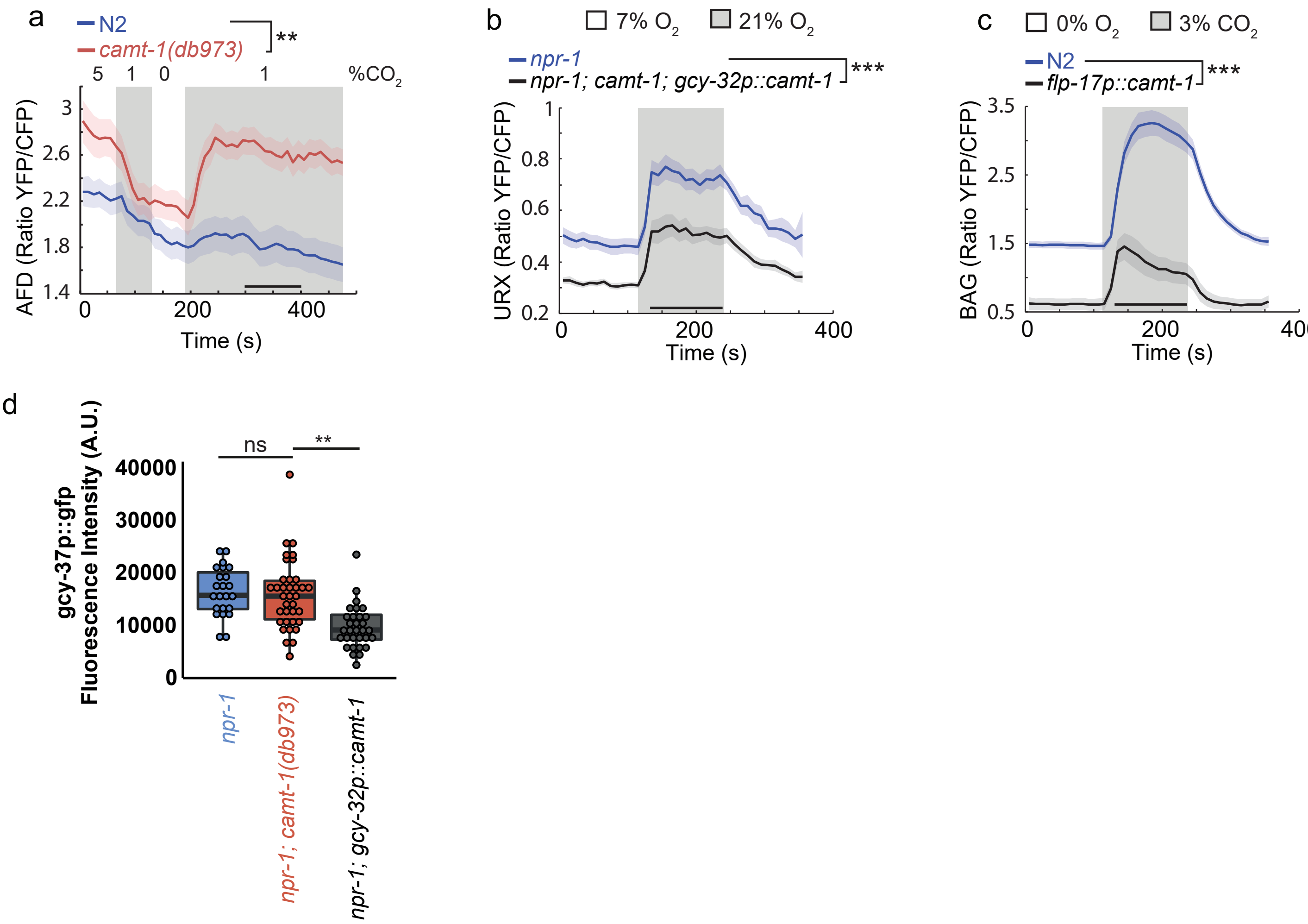

Extended Data Figure 4

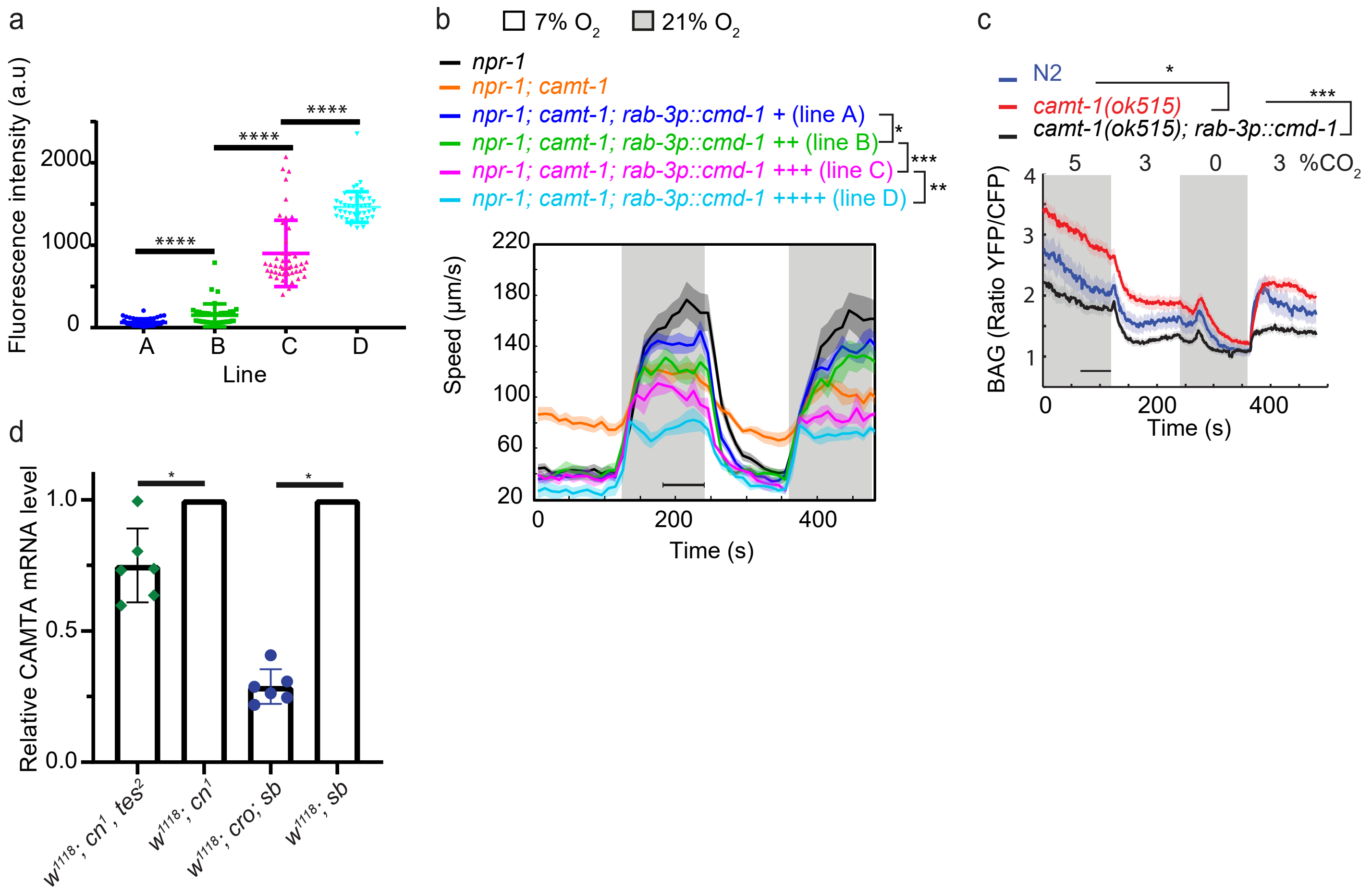

Extended Data Figure 5

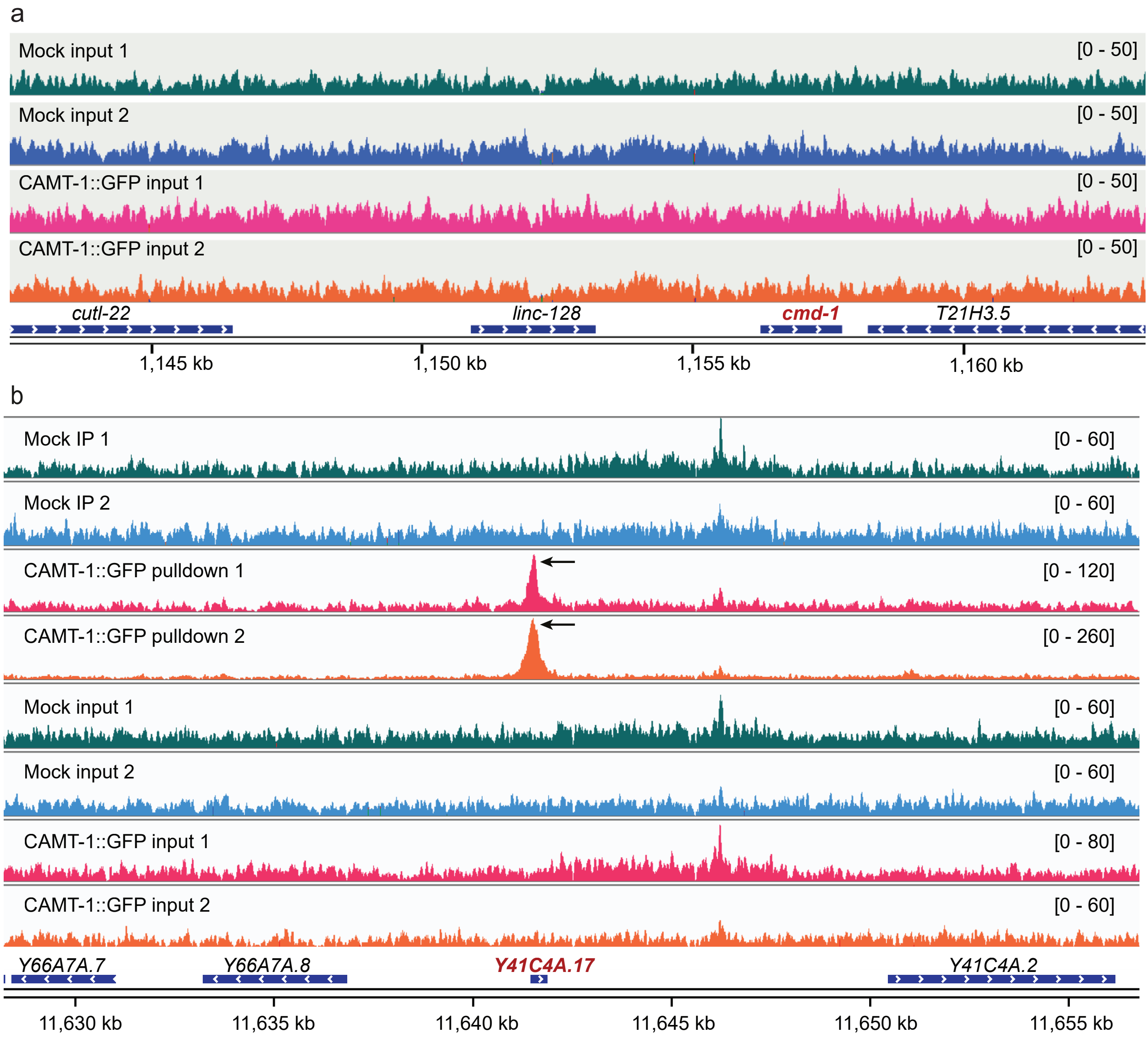
